# Comparative Transcriptomics of Human Breast Tissue Suggests Conserved Epithelial Secretory Programs and Tissue-Associated Regulatory Specialization

**DOI:** 10.64898/2025.12.19.695381

**Authors:** Marie Saitou

## Abstract

The mammary gland represents a defining evolutionary innovation of mammals, which is an epithelial tissue with specialized secretory functions that are regulated by hormonal and transcriptional processes. While mammary differentiation has been extensively studied, its transcriptional relationship to other epithelial tissues with secretory activity has been less explored in an evolutionary context.

Here, we used bulk RNA-seq data from the GTEx project to examine gene expression patterns in human breast tissue alongside a set of other tissues that contain epithelial secretory components, as well as non-glandular tissues. Using cross-tissue differential expression analyses, co-expression network analysis, and integration with single-cell expression references, we characterized patterns of gene expression that recur across multiple secretory tissues and those that show relative enrichment in breast tissue.

This analysis highlights three broad classes of expression patterns: genes with elevated expression across several secretory tissues, genes showing preferential expression in breast tissue, and genes displaying sex-associated differences in breast samples. These gene sets are associated with distinct functional annotations, including epithelial structure, metabolic and hormone-responsive processes, and immune-related functions.

Comparative analyses of gene conservation indicate that many genes preferentially expressed in breast tissue are evolutionarily conserved across vertebrates, consistent with the reuse of pre-existing gene repertoires in the mammalian lineage in tissue-specific contexts. Together, these results support a model in which breast tissue identity reflects the regulatory re-weighting and integration of conserved epithelial secretory modules rather than lineage-specific gene innovation. By placing human breast tissue gene expression in a comparative and evolutionary context, this study provides a molecular framework for understanding how complex organ-level traits arise through regulatory modularity.

**Significance:** The mammary gland is a defining feature of mammals, yet it remains unclear how its gene activity relates to that of other tissues with similar secretory functions. By comparing gene expression in human breast tissue with other epithelial secretory tissues, this study shows that much of breast tissue gene expression is shared with a common epithelial program, while a smaller set of genes is associated with breast-specific metabolic and hormonal functions. These findings help clarify how mammary gland specialization arises in mammalian lineages from ancestral genetic materials rather than from entirely new gene emergences, providing a framework for understanding the evolution of new organs.

## Introduction

The mammary gland, a hallmark trait of mammals, plays an essential role in mammalian offspring survival providing nutrition and immunity through milk (Peaker 2002; Hassiotou & Geddes 2013; Pond 1977). The mammary gland is a secretory tissue that undergoes coordinated changes in growth, differentiation, and function from embryogenesis through adulthood and lactation (Medina 1996). Its development involves interactions among epithelial, stromal, and immune compartments, regulated by hormonal signaling and transcriptional control mechanisms (Kaur et al. 2021; Rauner & Kuperwasser 2021; Dawson & Visvader 2021; Slepicka et al. 2021). Previous studies have described cellular hierarchies, stem and progenitor populations, and endocrine regulation associated with mammary morphogenesis and milk production (Chen et al. 2017; Slepicka et al. 2021; Tharmapalan et al. 2019).

From an evolutionary perspective, the mammary gland represents a major organ-level innovation whose origin predates the diversification of extant mammalian lineages. Classical studies based on comparative morphology and histology, proposed that mammary glands originated through the progressive modification of a single ancestral integumentary gland, typically hypothesized to be a sweat, sebaceous, or hair-associated gland (Gegenbauer 2017; Bresslau & Hill 1920; Blackburn 1991; Oftedal & Dhouailly 2013; Moll et al. 2008; Picardo et al. 2009). These models provided an important anatomical framework for understanding mammary gland origins.

In parallel, from the molecular evolution angle, comparative and phylogenetic studies have characterized a set of prominent milk-associated proteins and hormonal components across mammals, showing that broadly similar components are present across mammalian lineages(Sharp et al. 2014; Enjapoori et al. 2014; Kawasaki 2018). Such studies have been successful in describing the evolutionary histories and molecular properties of individual components of lactation. However, these approaches do not address how such components are transcriptionally organized into tissue-level gene expression programs, nor how these programs relate to those of other epithelial tissues.

As a result, a key gap remains in our understanding of mammary gland evolution: how gene expression patterns observed in breast tissue relate, at the transcriptome-wide level, to those of other epithelial tissues with secretory activity. Anatomical and functional comparisons indicate that the mammary gland shares features with other tissues containing epithelial secretory components, pancreas, stomach, and skin (McManaman et al. 2006). These tissues differ substantially in physiological roles and regulatory organization (Khan et al. 2022), but provide a context in which common and tissue-associated expression patterns can be examined. Comparing transcriptional profiles across such tissues offers a way to situate breast tissue gene expression within a broader landscape of epithelial biology (Brückner & Parker 2020).

Many transcriptomic and single-cell studies have focused on individual tissues, providing detailed descriptions of cellular composition and lineage organization within specific glands. In the mammary gland, single-cell RNA sequencing of tissue and milk has characterized epithelial and immune cell populations across developmental and physiological states (Li et al. 2020; Twigger et al. 2022). Comparable transcriptomic analyses have been performed for other secretory organs, including the salivary gland(Saitou et al. 2020; Gao et al. 2018; Hauser et al. 2020; Altrieth et al. 2024), pancreas (Ku et al. 2012; Altrieth et al. 2024; Bailey et al. 2016), stomach and intestinal glands (Huang et al. 2025; Öling et al. 2024)(Elmentaite et al. 2021), and skin tissues (Inoue et al. 2022).

However, these studies have largely addressed tissue-specific questions, and comparatively few analyses have examined how breast tissue gene expression relates to that of other epithelial secretory tissues within a unified comparative framework. To address this gap, we examined transcriptomic patterns in human breast tissue in comparison with multiple epithelial tissues containing secretory components, with the aim of distinguishing gene expression features that are broadly shared from those that show relative enrichment in breast tissue.

## Materials and Methods

### Dataset Selection and Tissue Group Definition

Bulk RNA-seq expression data were obtained from the GTEx v10 dataset. Tissues were categorized into two analysis groups for comparative purposes, referred to here as tissues containing epithelial secretory components and non-glandular tissues, for differential expression analyses.

The epithelial secretory tissue group comprised six tissues characterized by epithelial origin and secretory function: Breast – Mammary Tissue, Minor Salivary Gland, Skin – Not Sun Exposed (Suprapubic), Pancreas, Stomach, and Small Intestine – Terminal Ileum. Breast, salivary gland, and skin were included due to their abundant exocrine acinar or ductal structures (Pia-Foschini et al. 2003; Wang et al. 2017), despite differences in secreted products. Pancreas, stomach, and small intestine represent classical exocrine organs involved in digestive enzyme and fluid secretion. GTEx breast samples are derived from adult donors, and pregnancy or lactation status is not publicly available. Accordingly, breast samples are treated as representing adult, non-lactational breast tissue.

The non-glandular group consisted of six tissues used as comparative reference tissues lacking prominent epithelial secretory structures: Muscle – Skeletal, Heart – Left Ventricle, Artery – Tibial, Esophagus – Muscularis, Adipose – Subcutaneous, and Adipose – Visceral (Omentum).

#### Data Filtering

Samples were filtered using GTEx-provided quality metrics. Inclusion required RNA Integrity Number (SMRIN) ≥ 6.0, mapping rate (SMMAPRT) ≥ 0.8, sequencing efficiency (SMEXPEFF) ≥ 0.6, at least 50 million total reads (SMRDTTL), and 50 million uniquely mapped non-duplicate reads (SMMPPDUN) Samples were required to express at least 10,000 genes (SMGNSDTC).

Samples with high duplicate rates (SMDPMPRT > 0.6), elevated base mismatch rates (SMBSMMRT > 0.01), excessive chimeric reads (SMCHMRT > 0.015), UniVec contamination (SMUVCRT > 0.1), or extreme 3′ bias (SM3PBMN outside 0.4–0.75) were excluded. A minimum estimated library complexity of 30 million unique fragments (SMESTLBS) was required. Samples missing key metadata required for modeling were excluded. After filtering, samples from twelve tissue types were retained.

#### Covariates and Sample Metadata

Sample-level biological covariates included sex, age category, and Hardy death classification. The variable *SEX* is coded as 1 for male and 2 for female, while *AGE* corresponds to decade intervals (e.g., 20–29, 30–39, etc.). The *DTHHRDY* score describes the mode of death on a five-level Hardy scale, which serves as a proxy for postmortem condition.

Technical covariates included collection center (SMCENTER), nucleic acid batch (SMNABTCH), gene expression batch (SMGEBTCH), analyte type (ANALYTE_TYPE), and storage condition (SMAFRZE). Sequencing and RNA quality metrics used either for filtering or modeling included SMRIN, SMTSISCH, SMTSPAX, SMRDTTL, SMMAPRT, SMMPPDUN, SMEXPEFF, SMEXNCRT, SMRRNART, SMCHMRT, and SM3PBMN.

For differential expression modeling, SEX, AGE, SMCENTER, and SMEXPEFF were included as fixed effects. Samples missing any of these covariates were excluded prior to normalization and statistical analysis.

#### Differential Expression Analysis

Raw RNA-seq read count data from thirteen human tissues were obtained from the GTEx v10 dataset. Count matrices were imported directly from the compressed GTEx `.gct.gz` files, with sample columns matched to the curated metadata. The combined matrix was then used to create *DGEList* objects in *edgeR (Robinson et al. 2010)*, and lowly expressed genes were excluded using the `filterByExpr()` function with the relevant grouping factor. Counts were normalized by the trimmed mean of M-values (TMM) method to correct for differences in library size and sequencing depth.

Differential expression analyses were performed under several contrasts. In the first analysis, tissues containing epithelial secretory components were compared to non-glandular tissues to identify genes showing elevated expression across this tissue set. In the second analysis, breast tissue samples were contrasted against non-glandular tissues to identify genes showing relative enrichment in breast tissue within this comparison framework. A third comparison was carried out within breast tissue samples to detect sex-associated expression differences between male and female samples. For each analysis, a linear modeling framework was applied using the limma–voom pipeline. Coefficients corresponding to the main factor of interest were extracted from the fitted model. Genes were considered significantly differentially expressed when the absolute logL fold change exceeded one and the false discovery rate (Benjamini–Hochberg adjusted p-value) was below 0.001.

Differentially expressed genes were grouped into three expression categories defined operationally by their patterns across these contrasts. Genes upregulated in both breast tissue and other epithelial secretory tissues were classified as (1) “epithelial secretory–enriched genes”. Genes upregulated in breast tissue relative to both non-glandular tissues and other epithelial secretory tissues were defined as (2) “breast-enriched genes”. Finally, genes exhibiting higher expression in female relative to male breast samples were categorized as (3) “female-biased genes”.

#### Integration of Bulk RNA-seq data and Single-cell RNA-seq Data

Single-cell RNA-seq datasets of breast tissue and subcutaneous adipose tissue (Lazarescu et al. 2025; Reed et al. 2024) were downloaded at CELLxGENE Discover portal (CZI Cell Science Program et al. 2025) as reference expression resources to contextualize bulk RNA-seq data We used publicly available single-cell RNA-seq data from the Human Breast Cell Atlas (HBCA, global dataset, ∼803,000 cells), generated from normal adult human breast tissue and snRNA-seq of human subcutaneous adipose tissue - all cells (37,879 cells, GSE281356) (Lazarescu et al. 2025; Reed et al. 2024). In this study, we used subcutaneous adipose tissue as the comparator for reference breast adipose data because the adipose component of the human breast is anatomically and histologically classified as subcutaneous adipose tissue. The human breast is composed of glandular (mammary) and adipose tissue embedded in a stromal framework beneath the skin, with adipose tissue forming the major volume of the breast outside the glandular component(Nickell & Skelton 2005). Adipose tissue located beneath the skin is defined as subcutaneous adipose tissue, in contrast to visceral adipose tissue, which is localized around internal organs within the abdominal cavity (Ibrahim 2010).

Cell-type composition of bulk mammary gland RNA-seq data was estimated using a reference-based deconvolution approach implemented in the DeconRNA-seq (Gong & Szustakowski 2013) package (Bioconductor). Reference expression matrices were derived from single-cell RNA-seq datasets of human mammary and subcutaneous adipose tissues (Lazarescu et al. 2025; Reed et al. 2024). For each dataset, mean expression profiles were calculated for every annotated cell type, with genes as rows and cell types as columns.

From the resulting deconvolution output, adipocyte-related fractions were obtained by summing columns annotated as “adipocyte,” “mesenchymal stem cell of adipose tissue,” or similar lipid- associated terms. These estimated fractions were then integrated with GTEx sample metadata containing sex and age information. Age was converted to numeric values where possible and categorized into five intervals (≤30, 31–40, 41–50, 51–60, and ≥61 years). While pregnancy/lactation status was not available in the sample metadata, among female donors, samples from individuals aged 39 years or younger accounted for approximately 17% of the dataset, with the majority derived from donors aged 40 years and above.

### Single-cell Expression Profiling and Gene Clustering

Single-cell reference data were processed in R using the *zellkonverter (*https://bioconductor.org/packages/zellkonverter). and *DelayedArray* (https://code.bioconductor.org/browse/DelayedArray/) packages to enable efficient handling of large HDF5-backed matrices. Annotated expression data were imported from the `.h5ad` file, and cells were grouped according to their annotated cell type. To focus on biologically relevant features, only genes previously identified as differentially expressed were retained. Investigation on single-cell data was restricted to genes identified in bulk differential expression analyses, in order to examine how bulk-defined gene sets map onto reference cell populations.

For each annotated cell type, we calculated the detected fraction, defined as the proportion of cells in which a given gene was expressed (nonzero counts). This metric was used as comparative measure of gene activity across cell populations. Detected-fraction matrices were subjected to hierarchical clustering (Euclidean distance, complete linkage) to group genes with similar patterns across annotated cell types. Clustering was used to summarize broad patterns of cell-type association. Each dendrogram was divided into three clusters, and functional enrichment analyses were performed for each cluster using *g:Profiler* (*gprofiler2* package (Kolberg et al. 2020)) with Gene Ontology (BP, MF, CC), KEGG, and Reactome databases. Enrichment significance was assessed using the false discovery rate (FDR) correction.

#### Evolutionary Conservation Analysis

To assess patterns of evolutionary conservation among gene sets defined by breast-associated differential expression, we compiled a list of human genes differentially expressed in breast tissue and other epithelial secretory tissues and examined their orthologous relationships across representative vertebrate lineages using Ensembl BioMart (release ≥111). The ortholog presence matrix included 13 species covering major vertebrate clades: Eutheria (*mouse, Mus musculus; rat, Rattus norvegicus; rabbit, Oryctolagus cuniculus; pig, Sus scrofa; cattle, Bos taurus; goat, Capra hircus; sheep, Ovis aries*), Marsupialia (koala, *Phascolarctos cinereus*), Monotremata (platypus, *Ornithorhynchus anatinus*), Aves (chicken, *Gallus gallus*), Amphibia (*Xenopus tropicalis*), and Teleostei (zebrafish, *Danio rerio*; medaka, *Oryzias latipes*).

For each human Ensembl gene ID, ortholog presence (1) or absence (0) was determined in each species, resulting in a binary matrix that retained the original annotation of breast-enriched, epithelial secretory–enriched, and sex-associated differential expression categories. Genes were grouped according to their human tissue–based expression category, and the proportion of genes possessing identifiable orthologs in each species was calculated for each category, using the number of human genes in that category as the denominator. This analysis was used to compare relative levels of evolutionary conservation across expression-defined gene sets, rather than to infer the evolutionary origin of specific tissues or traits.

### Weighted gene co-expression network analysis (WGCNA) and cross-tissue module comparison

Normalized gene expression counts from the GTEx dataset were used to construct tissue-specific co-expression networks using the WGCNA package (v1.72). For breast tissue, cell-type composition effects estimated by DeconRNA-seq were regressed out to reduce variability associated with major non-epithelial components, specifically endothelial and macrophage fractions, using removeBatchEffect in the limma package.

Normalized gene expression counts from the GTEx dataset were used to construct tissue-specific co-expression networks using the WGCNA package (https://cran.r-project.org/web/packages/WGCNA/index.html) (v1.72). For the breast tissue, cell-type composition effects estimated by DeconRNA-seq were regressed out to reduce variability associated with endothelial and macrophage fractions, using removeBatchEffect in the limma package. using `removeBatchEffect` in the limma package (Ritchie et al. 2015). Log-transformed CPM values were filtered to retain genes expressed in at least 20% of samples, and a signed network was built with bicor correlation and dynamic tree cutting (minimum module size = 120, merge cut height = 0.35). Modules were labeled in descending order of size (Module 1, Module 2,…), and module eigengenes (MEs) were extracted for downstream analyses.

To evaluate similarity of co-expression structure across tissues, module assignments from the mammary gland were compared with those from other GTEx tissues. For each tissue, Jaccard similarity was computed between all pairs of mammary and tissue modules based on shared gene memberships. For each breast module, the maximum Jaccard index across modules in a given tissue was retained as a summary measure of overlap (“best-per-tissue” score). Permutation tests (n = 2000) were performed by shuffling gene labels in the comparison tissue while preserving module sizes, yielding empirical p-values and FDR-corrected significance estimates.

#### Use of Large Language Model

We used ChatGPT (version 5.2, OpenAI) to assist with code refactoring, language editing, and proofreading. All scientific interpretations, analyses, and conclusions are the responsibility of the authors.

## Results

### Comparative transcriptomic analyses reveal patterns of breast-tissue-associated gene expression

Principal component analysis (PCA) was performed using the top 2,000 most variable genes across 12 tissue types. The first two components explained 37.6% (PC1) and 19.7% (PC2) of the variance, respectively. Samples from tissues containing epithelial secretory components, including breast, salivary gland, pancreas, stomach, small intestine, and skin, formed a coherent cluster that was separated from non-glandular tissues such as muscle, heart, esophagus, and adipose. Breast tissue samples occupied a position intermediate between adipose tissue and other epithelial secretory tissues, while remaining closer to the latter in PCA space. Breast, salivary gland, and pancreas samples showed relative proximity along the first two principal components (**Figure 1**).

**Figure 1.**
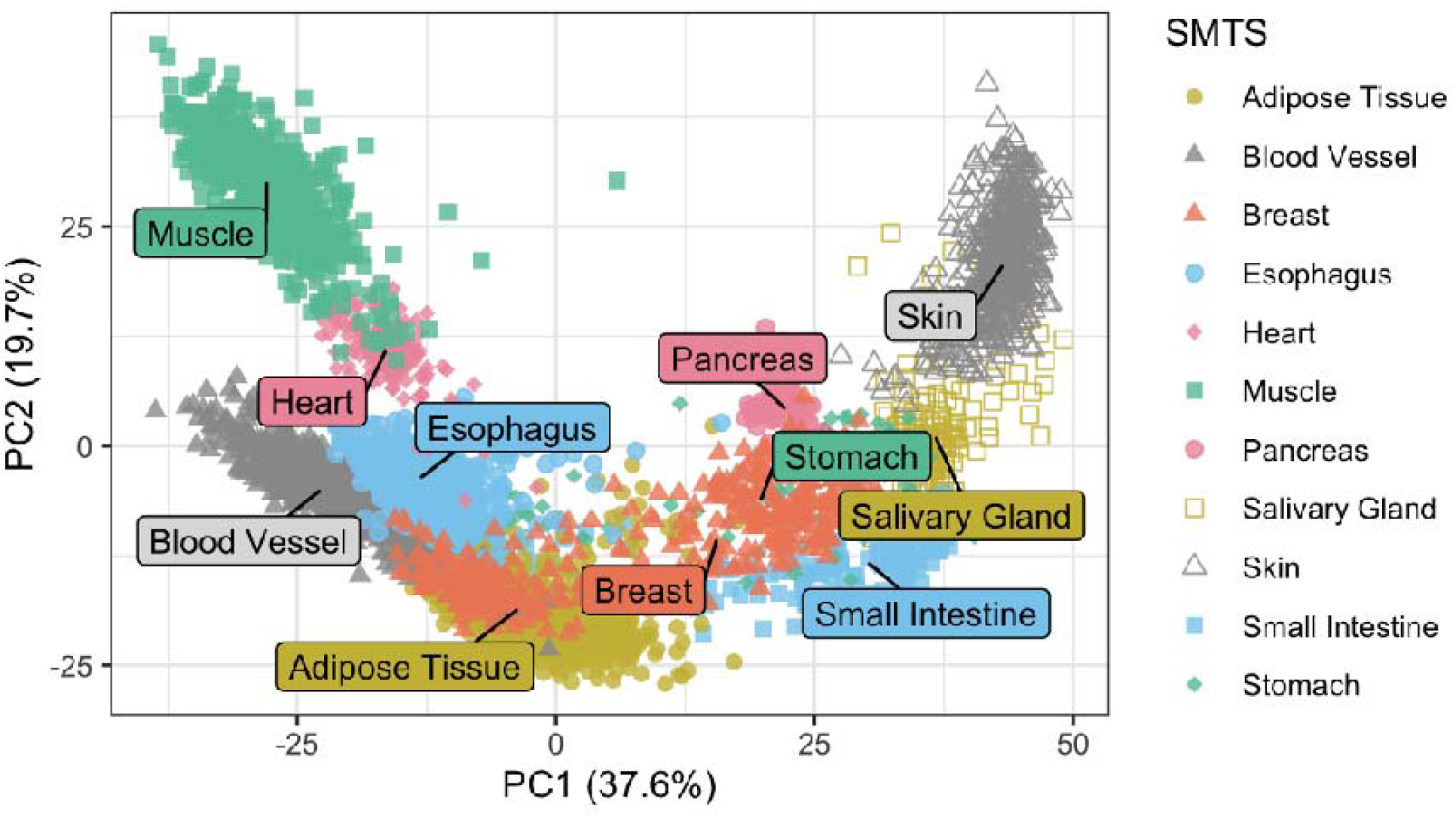
Principal component analysis (PCA) of tissue transcriptomes. Principal component analysis (PCA) was performed using the 2,000 genes with the highest expression variance across tissues, to summarize global patterns of transcriptome variation across samples. Each point represents a tissue sample, and tissues are colored according to their anatomical source (legend, right). Tissues containing epithelial secretory components (breast, salivary gland, pancreas, stomach, small intestine, skin) cluster separately from non-glandular tissues (muscle, heart, blood vessel, esophagus, adipose). The x-axis (PC1) and y-axis (PC2) show the first and second principal components, which together explain the largest proportions of total expression variance (37.6% and 19.7%, respectively). Samples that are closer together in this plot have more similar global expression profiles, whereas those farther apart differ more strongly in their transcriptome composition. Colored clusters correspond to distinct tissue types. Labels indicate the centroids of major tissue groups.

We next compared gene expression profiles across three expression categories defined by differential expression patterns across tissues and sex. Differentially expressed genes were grouped into three categories based on their patterns of relative enrichment. Genes upregulated in both breast tissue and other epithelial secretory tissues were classified as (1) “epithelial secretory–enriched genes” (711 genes). Genes showing higher expression in breast tissue relative to both non-glandular tissues and other epithelial secretory tissues were defined as (2) “breast-enriched genes” (189 genes). Finally, genes exhibiting higher expression in female relative to male breast samples were categorized as (3) “female-biased genes” (617 genes) (**Figure 2A**; **Table S1**).

**Figure 2.**
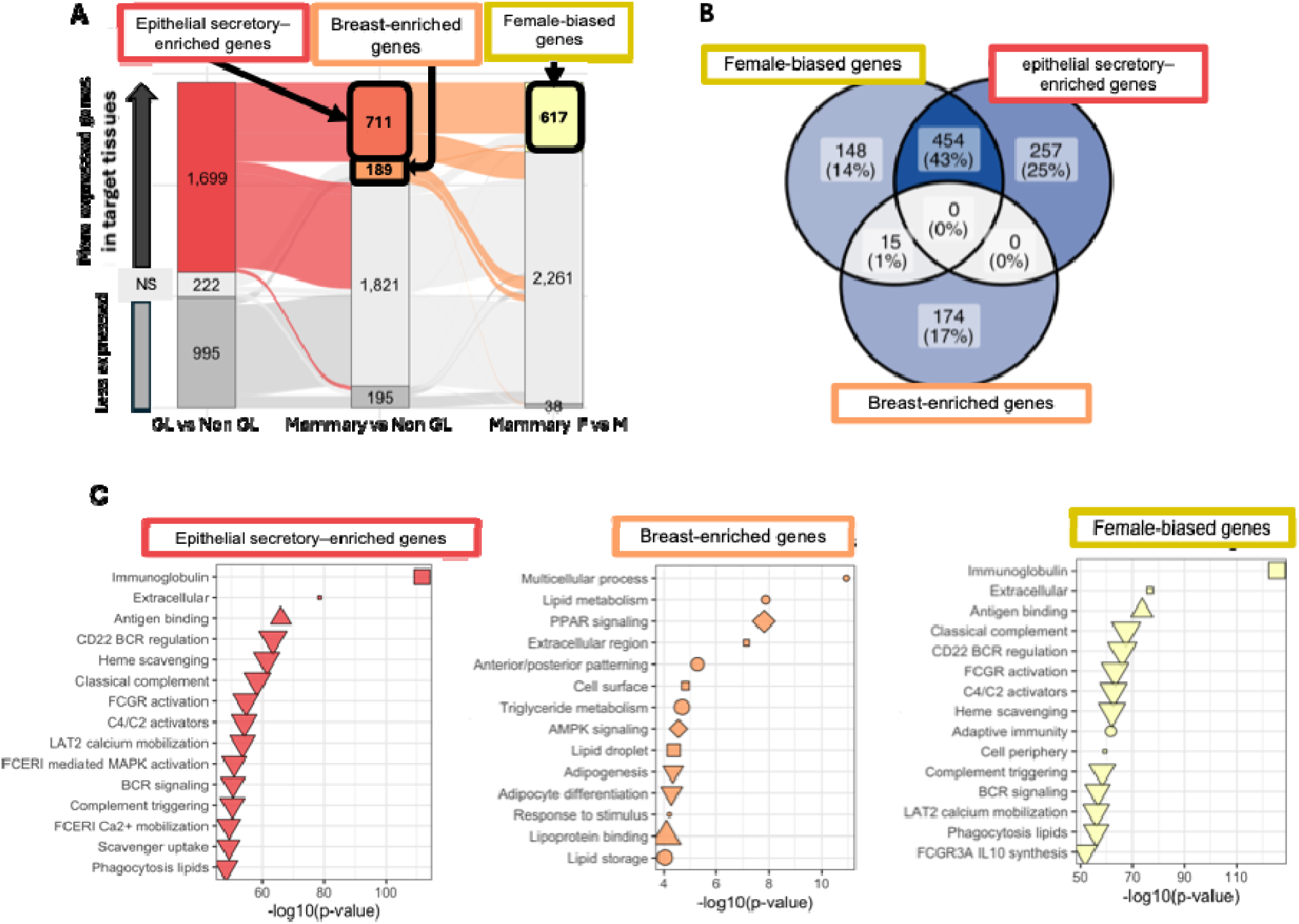
Differentially expressed gene categories across comparisons. (A) Sankey diagram summarizing the number of differentially expressed genes (DEGs) identified in three pairwise comparisons: (i) tissues containing epithelial secretory components versus non-glandular tissues, (ii) breast tissue versus non-glandular tissues, and (iii) female versus male breast tissue samples. Bars indicate the number of upregulated (colored), downregulated (grey), and non-significant (NS; white) genes. Flows illustrate overlaps of gene categories across comparisons. (B) Venn diagram illustrating the overlap among three categories of upregulated genes: epithelial secretory–enriched genes, breast-enriched genes, and female-biased genes. Numbers and percentages indicate the proportion of genes within each intersection. (C) Gene Ontology (GO) and pathway enrichment analyses performed using g:Profiler for upregulated genes in the three DEG categories.

We observed substantial overlap between epithelial secretory–enriched genes and female-biased genes, whereas breast-enriched genes showed limited overlap with either category (**Figure 2B**). Functional enrichment analysis further showed that these gene sets were associated with distinct functional annotations (**Figure 2C; Table S2**). epithelial secretory–enriched genes were predominantly enriched for immune-related and extracellular functions, breast-enriched genes were enriched for metabolic and endocrine pathways, and female-biased genes showed strong enrichment for immune and complement-associated processes.

In the “epithelial secretory–enriched genes” group, we identified a cluster of genes associated with epithelial structure, secretion, and ion transport. This set included *DSP* (desmoplakin), *CDH1* (E-cadherin), *GRHL2*, and *BARX2*, which regulate epithelial integrity and branching morphogenesis, as well as several solute carriers *(SLC13A2, SLC5A1, SLC15A1*) and secretory enzymes (*PRSS8, PRSS22, SULT2B1*). In contrast, the “breast-enriched genes” group was characterized by genes associated with hormonal signaling, metabolic specialization, and epithelial differentiation. This set included hormone- and metabolite-responsive receptors such as *OXTR* (oxytocin receptor) and *HCAR1* (lactate receptor), together with the developmental regulator *TBX3* and the mammary-enriched long non-coding RNA *LINC00993*. Additional genes in this category comprised regulators of lipid metabolism, immune-associated processes, and cellular stress responses (e.g. *CREB3L4, STC2, CMTM7, SAA2,* and *SAA4*). The group further contained multiple transcription factors and structural or metabolic genes linked to epithelial organization and function, including *TWIST1, HOXD9, TRPS1, CLPSL1, PEMT,* and *SLC5A6*. Finally, the “Female-biased genes” category was dominated by immunoglobulin genes, including IGHV, IGKJ, IGLJ, and IGKC family members, together with POU2AF1 and CD27.

#### Single-cell mapping and bulk deconvolution provide a cellular context for breast tissue gene expression patterns

Bulk RNA-seq data from GTEx breast tissue were deconvoluted using single-cell breast and adipose reference profiles from the CELLxGENE Discover portal to obtain descriptive estimates of cell-type composition across sex and age groups. Across all age groups and in both sexes, breast tissue samples contained a substantial adipose-associated fraction, accounting for approximately half of the inferred cellular composition (**Figure S1**). While the relative proportion of adipose-associated cells varied across samples, the overall composition was similar across sex and age categories.

The remaining fraction consisted primarily of annotated epithelial and stromal-associated cell populations, including basal–myoepithelial and perivascular cells, with smaller contributions from macrophage and endothelial compartments. Given the age distribution of GTEx breast tissue donors, which is skewed toward middle-aged and older individuals, the inferred cellular composition is consistent with adult, non-lactational breast tissue samples. This compositional profile parallels the relative positioning of breast tissue samples between adipose and other epithelial secretory tissues in the global PCA (**Figure 1A**).

To determine how each differential expression category is distributed across mammary cell populations, we examined the cellular expression patterns of genes belonging to the three defined groups: “epithelial secretory–enriched genes”, “breast-enriched genes”, and “female-biased genes”. For each gene, we calculated the detected fraction, defined as the proportion of cells within each annotated mammary cell type in which the gene was expressed (**Figure 3**).

**Figure 3.**
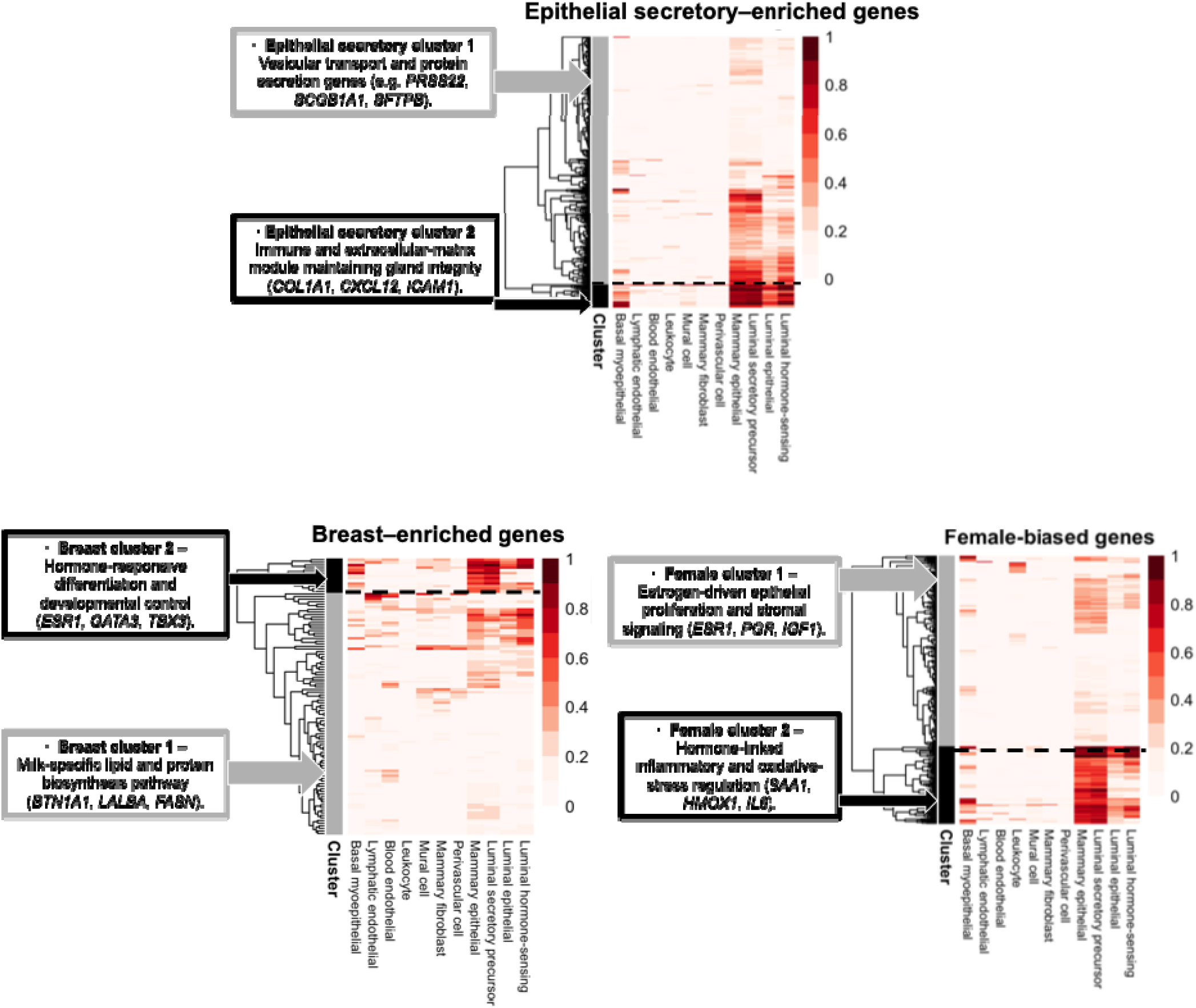
Single-cell expression patterns of upregulated gene sets. Heatmaps showing the proportion of single-cell populations expressing upregulated genes in each DEG category: (i) genes upregulated across glandular tissues, (ii) genes upregulated only in mammary gland, and (iii) genes upregulated in female versus male mammary gland. Cell types include basal myoepithelial cells, luminal epithelial subtypes (hormone-sensing, secretory precursor, differentiated luminal), fibroblasts, endothelial cells (blood and lymphatic), mural cells, perivascular cells, and leukocytes. Expression levels are represented by scaled proportions of expressing cells (0–1), with rows corresponding to genes and columns to cell types.

Within the “epithelial secretory–enriched genes” category, hierarchical clustering separated genes into two major clusters reflecting distinct single-cell detection patterns. One cluster comprised genes broadly detected across multiple mammary epithelial populations, including genes associated with vesicular transport and protein secretion, such as *PRSS22*. A second group showed higher detected fractions in annotated epithelial, secretory precursor, and hormone-sensing cell populations. These genes are associated with extracellular matrix and immune-related components, including *COL1A1*, *CXCL12*, and *ICAM1*.

For “breast-enriched genes”, cluster 1 showed a broad and heterogeneous expression pattern and included genes linked to mammary differentiation and metabolic specialization such as *BTN1A1*, and *LALBA*. Cluster 2 genes were expressed in mammary epithelial and luminal secretory precursor cells, and encompassed genes associated with hormone-responsive differentiation and developmental control, such as *ESR1, GATA3*, and *TBX3*.

Genes in the “female-biased genes” category also segregated into two major cellular expression patterns. Cluster 1 showed broad expression, and included estrogen-driven epithelial proliferation genes and stromal signaling genes, including *ESR1, PGR* and *IGF1*. Cluster 2 genes were expressed in mammary epithelial and luminal secretory precursor cells, and associated with hormone-linked inflammatory and oxidative stress regulation, including *SAA1, HMOX1* and *IL6*.

#### Comparative genomics analyses compare patterns of evolutionary conservation across expression-defined breast gene categories

To examine patterns of evolutionary conservation among gene sets defined by tissue-associated expression, we compared the evolutionary conservation of the three gene categories identified in this study. Specifically, for genes classified as “epithelial secretory–enriched genes”, “breast-enriched genes”, and “female-biased genes”, we assessed the presence or absence of identifiable orthologs across a set of vertebrate species, including both mammalian and non-mammalian lineages spanning monotremes, birds, amphibians, and teleost fish (**Table S3**).

Comparison of ortholog retention ratios indicated differences among the three gene categories (Kruskal–Wallis test, χ² = 32.25, df = 2, p = 9.9 × 10LL; **Figure 4)**. Pairwise comparisons showed that genes classified as “breast-enriched” exhibited higher ortholog retention across the surveyed vertebrate species than genes classified as “epithelial secretory–enriched” (Wilcoxon test, p = 3.5 × 10LL) or “female-biased” (Wilcoxon test, p = 6.1 × 10LL).

**Figure 4.**
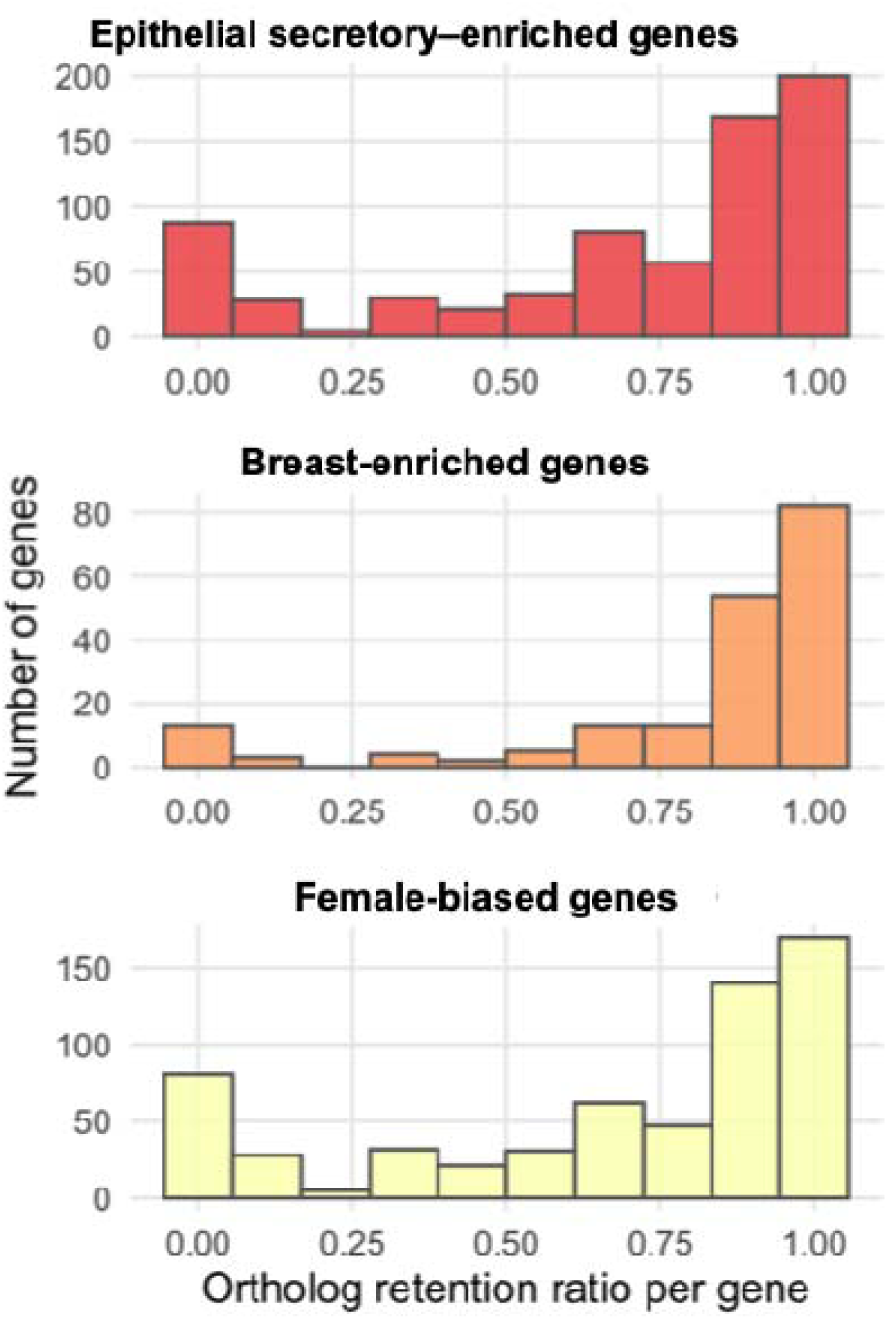
Evolutionary conservation of mammary gene categories across vertebrate species. For each gene category, the presence or absence of orthologs of human genes, (genes in other species that descend from a common ancestral gene), was examined across vertebrate species: Mouse, Rat, Cattle, Goat, Sheep, Pig, Rabbit, Koala, Platypus, Chicken, Xenopus tropicalis, Zebrafish, and Medaka. A gene’s ortholog retention ratio represents the fraction of these species in which a corresponding ortholog is detected (0 = no orthologs retained, 1 = orthologs present in all species). Histograms display the distribution of retention ratios for three categories: epithelial secretory–enriched genes (red), breast-enriched genes (orange), and Female-biased genes (yellow). The x-axis shows the ortholog retention ratio per gene, and the y-axis indicates the number of genes in each bin.

#### Mammary co-expression modules and their overlap with other glandular tissues

To examine patterns of gene co-expression in breast tissue and their similarity to those observed in other tissues, we performed weighted gene co-expression network analysis (WGCNA). WGCNA of breast tissue transcriptomes identified 14 co-expression modules, ranging in size from 232 to 2,429 genes (median ≈ 540 genes). The two largest modules (Modules 1 and 2) contained more than 1,000 genes each, whereas the remaining modules comprised between 200 and 900 genes.

To assess similarity of co-expression patterns across tissues, we calculated pairwise Jaccard indices between breast-derived modules and modules derived from 12 other GTEx tissues (**Figure S2**). Overall overlap between breast modules and those from other tissues was low, with median Jaccard indices below 0.1 and no detectable difference between tissues containing epithelial secretory components and non-glandular tissues (Wilcoxon test, p > 0.05). Among the identified modules, breast Module 2 showed the highest and most recurrent overlap with modules from several tissues containing epithelial secretory components, including salivary gland, pancreas, stomach, and skin (**Figure 5A**).

**Figure 5.**
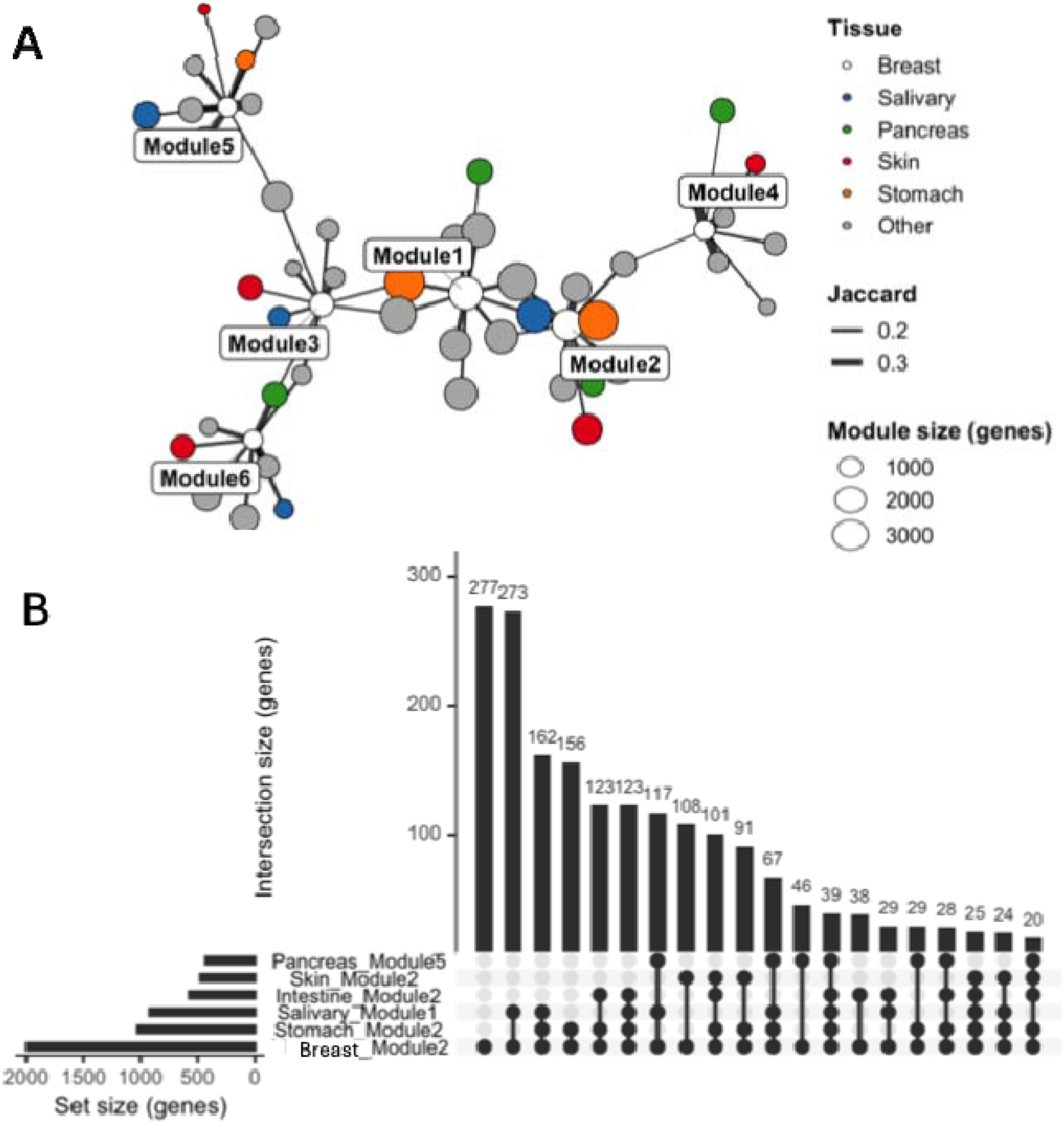
Cross-tissue network comparison of mammary gene co-expression modules and their gene set overlaps. **(A)** Network visualization of breast-specific gene co-expression modules (Modules 1–6) and their best-matching modules identified across other human tissues. Each node represents a gene co-expression module, defined as a cluster of genes that show highly correlated expression patterns within a given tissue, derived from tissue-resolved network analysis. Node size corresponds to the number of genes included in each module. Node color indicates the tissue of origin, with white nodes representing breast modules and colored nodes representing modules from other glandular tissues. Edge thickness is proportional to the Jaccard index value, such that thicker lines indicate stronger gene overlap between modules. **(B)** UpSet plot summarizing gene set intersections between breast modules and their top-matching modules from other tissues. Each filled dot in the bottom matrix represents the inclusion of a particular module in a given intersection set, and connecting lines indicate combinations of modules that share overlapping genes. The vertical bars above the matrix show the number of genes (intersection size) for each specific module combination, while the horizontal bars on the left indicate the total number of genes (set size) contained in each module.

To describe the gene composition underlying these overlaps, we examined gene subsets shared between breast Module 2 and one or more comparison tissues. For each comparison tissue, only the module with the highest overlap with breast Module 2 was considered (**Figure 5B**). Genes unique to the breast module were associated with pathways related to lipid metabolism and mitochondrial processes, including fatty acid biosynthesis, β-oxidation, and PPAR signaling, as well as hormone-associated signaling pathways. Gene sets shared between breast tissue and salivary gland or pancreas were associated with pathways related to vesicle trafficking, Golgi–ER processing, glycosylation, and exocytosis. Gene sets shared across breast tissue, salivary gland, and stomach were associated with extracellular matrix organization (**Table S4**).

## Discussion

Our analyses indicate that the breast tissue shares transcriptional features with other epithelial tissues containing secretory components in humans. Genes classified as epithelial secretory–enriched were expressed in the mammary gland as well as in salivary gland, pancreas, stomach, and skin, and were primarily associated with epithelial structure, secretory processes, extracellular matrix organization, and immune-related functions. This pattern indicates that a substantial fraction of breast tissue gene expression reflects transcriptional programs that are not unique to the mammary gland, but are broadly deployed across epithelial secretory tissues. The epithelial secretory–enriched genes identified here are consistent with patterns reported in prior transcriptomic analyses in mouse, which showed that exocrine organs such as the salivary gland, pancreas, and skin exhibit broadly similar epithelial gene expression profiles at the global level (Gluck et al. 2016). Single-cell reference mapping further indicated that many of these shared expression patterns are detectable in annotated epithelial cell populations, suggesting that common glandular features are largely carried by epithelial cell types with secretory or barrier-related functions.

Genes classified as glandular-enriched included regulators of epithelial adhesion and associated with breast cancer, such as *CDH1* (Zhang et al. 2023; Dopeso et al. 2024), transcription factors implicated in epithelial differentiation including *GRHL2(Wang et al. 2023; Kumegawa et al. 2022)* and *BARX2(Yanan Sun et al. 2024)*. These genes have also been extensively studied in the context of breast cancer, where they are linked to epithelial identity and disease-associated transcriptional programs. Their enrichment across multiple epithelial secretory tissues suggests that such regulators form part of a shared epithelial gene expression core rather than representing mammary-specific innovations. Epithelial secretory–enriched genes also included solute carriers involved in epithelial transport, such as *SLC15A1*, which is well characterized for its duodenal-specific expression and role in peptide transport in the human upper gastrointestinal tract (Thwaites and Anderson, 2007; Hediger et al., 2013). Its presence among epithelial secretory–enriched genes is consistent with the use of established epithelial transport mechanisms across multiple tissues with absorptive or secretory functions.

In contrast, breast-enriched genes were enriched for regulators of hormonal responsiveness, metabolism, and epithelial differentiation. This group included hormone-related receptors such as *OXTR* (Reversi et al. 2005; Liu et al. 2020; Li et al. 2021, 2018) and *HCAR1(Mohammad Nezhady et al. 2023; Jin et al. 2022; Wagner et al. 2017)*, transcription factors involved in mammary development and lineage specification including *TBX3 (Douglas & Papaioannou 2013)*, *TRPS1(Cornelissen et al. 2020)*, and *TWIST1(Xu et al. 2017; Foubert et al. 2010)*, and metabolic regulators such as *PEMT* (Resseguie et al. 2007) and *SLC5A6(Montalbetti et al. 2014)*. The functional annotations of this gene set are consistent with known metabolic and endocrine features of breast tissue, particularly in relation to milk production. Several of these genes are known to respond to hormonal cues involved in mammary differentiation and secretory activation, and their expression patterns align with established roles of endocrine and metabolic regulation in mammary epithelial function (Li et al. 2018). These observations suggest that breast-enriched expression patterns are associated with metabolic and hormone-responsive processes layered onto a broader epithelial transcriptional background.

Genes exhibiting female-biased expression were dominated by B cell-related immunoglobulin genes, including *IGKC*, *IGHM*, *IGHV*, and *IGKJ* family members, which are well characterized as components of B cell and antibody-mediated immune responses (Hill et al. 2023)(Lycke et al. 2015). In addition, immune-associated regulators such as *POU2AF1* (Wenxiang Sun et al. 2024) and *CD27* (Jaeger-Ruckstuhl et al. 2024), which are primarily involved in T cell function and immune modulation, were also enriched. The observed female-biased expression of immune-related genes likely reflects differences in immune cell representation and activation states within breast tissue, in addition to sex-specific regulation of epithelial gene expression.

Taken together, these results are consistent with an interpretation in which breast tissue gene expression reflects a combination of expression patterns shared with other epithelial secretory tissues and a more limited set of features showing relative enrichment in breast tissue. Rather than indicating a wholly distinct mammary-specific transcriptional program or widespread lineage-specific gene innovation, the breast transcriptome appears to be structured around conserved epithelial-associated gene sets with tissue-associated modulation linked to metabolic and hormonal functions. Consistent with these observations, hormone-associated regulatory mechanisms have been proposed as key drivers of tissue-specific switch-like expression patterns (Aqil et al. 2025), providing a plausible framework for interpreting mammary-specific and female-biased gene programs identified in the present study. This organization is compatible with the co-expression analyses presented in this study, which show partial overlap between breast-derived co-expression modules and those identified in other epithelial secretory tissues.

Within this framework, breast-associated genes may be interpreted as components of conserved gene sets that are differentially weighted or regulated in breast tissue. One example is *STC2 (Stanniocalcin 2),* a mammary-specific gene in our analysis. Although stanniocalcins originated as calcium-regulatory hormones in non-mammalian vertebrates, STC2 is expressed in adult, non-lactational mammary tissue and its expression pattern is consistent with the reuse of an ancestral endocrine regulator in a breast-associated context (Hasilo et al. 2005).

From an evolutionary perspective, the patterns observed here are compatible with models in which tissue specialization arises through regulatory modification of conserved gene repertoires. Studies in both plants and animals have shown that novel organ morphologies often emerge through changes in the deployment, connectivity, or regulation of pre-existing gene modules rather than through the invention of new genes (e.g. “old genes, new functions”) (Rosin & Kramer 2009; Defoort et al. 2018; Pires & Dolan 2012; Uebbing et al. 2024). In this context, breast tissue can be viewed as assembling transcriptional features drawn from broadly conserved epithelial and secretory gene programs, together with regulatory modifications associated with mammary-specific metabolic, endocrine, and reproductive functions. This interpretation is consistent with the observed evolutionary conservation of many breast-associated genes across vertebrates, indicating that breast tissue specialization is associated with regulatory reorganization of conserved gene sets rather than lineage-specific gene invention.

Together, these findings situate breast tissue gene expression within a continuum of epithelial tissues containing secretory components, highlighting shared transcriptional features alongside tissue-associated expression patterns. Viewing breast gene expression through this comparative framework provides a basis for integrating transcriptomic observations across tissues, and may be applicable to the study of other epithelial organs that exhibit both shared molecular features and tissue-specific physiological specialization. Viewing breast gene expression through this comparative and evolutionary framework provides a basis for interpreting mammary gland identity as an outcome of regulatory modularity, and suggests that similar approaches may be informative for understanding the evolution of other epithelial organs that combine conserved molecular architectures with tissue-specific physiological functions.

## Supporting information

STables

SFigures

## Data Availability

All data and analysis scripts supporting this study are available at: https://github.com/mariesaitou/paper_2025-/tree/main/breast_comparative

The study utilized publicly accessible human transcriptomic datasets from the GTEx Consortium (v10) and the CZ CELLxGENE Discover portal, including the Human Breast Cell Atlas and human subcutaneous adipose tissue datasets (See **Materials and Methods**). No additional restrictions apply to data sharing.

## Acknowledgments

Computational analyses were performed using the High Performance Computing (HPC) cluster *Orion* at the Norwegian University of Life Sciences (NMBU). We thank Sabine Anne-Lie Ferneborg (NMBU, BIOVIT) for helpful comments and constructive feedback on the manuscript.

## Author Contributions

M.S. designed the study and conducted the bioinformatics analysis. G.K.S. interpreted glandular gene programs. Both authors contributed to writing the manuscript and approved the final version.

## Ethical Approval

This study utilized publicly available human transcriptomic datasets (GTEx and CellxGene). No new human or animal samples were collected for this work.

## Supplemental Material

**Table S1. Differentially expressed genes defining glandular-enriched, mammary-specific, and female-biased expression programs**

List of differentially expressed genes identified from GTEx RNA-seq analyses and classified into three expression categories: epithelial secretory–enriched genes, breast-enriched genes, and female-biased genes.

**Table S2. Functional enrichment of single-cell gene clusters based on detected fraction patterns**

Functional enrichment results from g:Profiler for gene clusters identified by hierarchical clustering of detected fraction profiles across mammary cell types. Enrichment is reported for Gene Ontology (BP, MF, CC), KEGG, and Reactome categories, with FDR-adjusted p-values.

**Table S3. Evolutionary conservation of expression-defined gene categories across vertebrates**

Presence or absence of orthologs for genes in each expression category across representative vertebrate species. Orthology was assessed using Ensembl BioMart, and conservation is reported as binary presence across species.

**Table S4. Functional enrichment of gene subsets from breast co-expression Module 2 shared with other epithelial secretory tissues**

g:Profiler enrichment results for gene subsets defined by overlap between breast co-expression Module 2 and modules from other tissues containing epithelial secretory components.

## Supplementary Figures

**Figure S1. Mean cell type composition in breast tissue by sex and age group**

Bulk RNA-seq data from GTEx breast tissue were deconvoluted using single-cell reference expression profiles to estimate relative cell-type composition. Stacked bar plots show mean fractional compositions averaged within each sex (male, female) and age group (10-year bins), with the y-axis normalized to 100%. Colors denote inferred cell populations: adipocyte / adaptive secretory precursor cell (A: Adipose), macrophages (A: Adipose), endothelial cells (A: Adipose), basal–myoepithelial cells (M: Mammary),perivascular cells (M: Mammary), and minor fractions classified as “Other cell types.” Single-cell RNA-seq reference datasets of breast tissue and subcutaneous adipose were downloaded at CELLxGENE Discover portal (CZI Cell Science Program et al. 2025).

**Figure S2. Cross-tissue overlap of breast co-expression modules.**

Heatmap showing Jaccard indices between breast tissue co-expression modules and modules derived from other GTEx tissues. For each comparison tissue, the Jaccard index represents the maximum overlap observed between a given breast module and any module from that tissue (“best-per-tissue” overlap). Rows correspond to comparison tissues, and columns correspond to breast co-expression modules. Tissues are annotated as tissues containing epithelial secretory components or non-glandular tissues based on the tissue grouping used in this study. Warmer colors indicate higher Jaccard similarity.

## References

Altrieth AL et al. 2024. Single-cell transcriptomic analysis of salivary gland endothelial cells. J. Dent. Res. 103:269–278.

Aqil A et al. 2025. Switch-like gene expression modulates disease risk. Nat. Commun. 16:5323.

Bailey P et al. 2016. Genomic analyses identify molecular subtypes of pancreatic cancer. Nature. 531:47–52.

Blackburn DG. 1991. Evolutionary origins of the mammary gland. Mamm. Rev. 21:81–96.

Bresslau E, Hill JP. 1920. The mammary apparatus of the mammalia : in the light of ontogenesis and phylogenesis /. Methuen & Co.,: London:

Brückner A, Parker J. 2020. Molecular evolution of gland cell types and chemical interactions in animals. J. Exp. Biol. 223:jeb211938.

Chen X, Liu Q, Song E. 2017. Mammary stem cells: angels or demons in mammary gland? Signal Transduct. Target. Ther. 2:16038.

Cornelissen LM et al. 2020. TRPS1 acts as a context-dependent regulator of mammary epithelial cell growth/differentiation and breast cancer development. Genes Dev. 34:179–193.

CZI Cell Science Program et al. 2025. CZ CELLxGENE Discover: a single-cell data platform for scalable exploration, analysis and modeling of aggregated data. Nucleic Acids Res. 53:D886–D900.

Dawson CA, Visvader JE. 2021. The cellular organization of the mammary gland: Insights from microscopy. J. Mammary Gland Biol. Neoplasia. 26:71–85.

Defoort J, Van de Peer Y, Vermeirssen V. 2018. Function, dynamics and evolution of network motif modules in integrated gene regulatory networks of worm and plant. Nucleic Acids Res. 46:6480–6503.

Dopeso H et al. 2024. Genomic and epigenomic basis of breast invasive lobular carcinomas lacking CDH1 genetic alterations. NPJ Precis. Oncol. 8:33.

Douglas NC, Papaioannou VE. 2013. The T-box transcription factors TBX2 and TBX3 in mammary gland development and breast cancer. J. Mammary Gland Biol. Neoplasia. 18:143–147.

Elmentaite R et al. 2021. Cells of the human intestinal tract mapped across space and time. Nature. 597:250–255.

Enjapoori AK et al. 2014. Monotreme lactation protein is highly expressed in monotreme milk and provides antimicrobial protection. Genome Biol. Evol. 6:2754–2773.

Foubert E, De Craene B, Berx G. 2010. Key signalling nodes in mammary gland development and cancer. The Snail1-Twist1 conspiracy in malignant breast cancer progression. Breast Cancer Res. 12:206.

Gao X, Oei MS, Ovitt CE, Sincan M, Melvin JE. 2018. Transcriptional profiling reveals gland-specific differential expression in the three major salivary glands of the adult mouse. Physiol. Genomics. 50:263–271.

Gegenbauer C. 2017. Zur Kenntniss Der Mammarorgane Der Monotremen. Hansebooks.

Gluck C et al. 2016. RNA-seq based transcriptomic map reveals new insights into mouse salivary gland development and maturation. BMC Genomics. 17:923.

Gong T, Szustakowski JD. 2013. DeconRNASeq: a statistical framework for deconvolution of heterogeneous tissue samples based on mRNA-Seq data. Bioinformatics. 29:1083–1085.

Hasilo CP et al. 2005. Nuclear targeting of stanniocalcin to mammary gland alveolar cells during pregnancy and lactation. Am. J. Physiol. Endocrinol. Metab. 289:E634–42.

Hassiotou F, Geddes D. 2013. Anatomy of the human mammary gland: Current status of knowledge. Clin. Anat. 26:29–48.

Hauser BR et al. 2020. Generation of a single-cell RNAseq atlas of Murine salivary gland development. iScience. 23:101838.

Hill L et al. 2023. Igh and Igk loci use different folding principles for V gene recombination due to distinct chromosomal architectures of pro-B and pre-B cells. Nat. Commun. 14:2316.

Huang RJ et al. 2025. A spatial transcriptomic signature of 26 genes resolved at single-cell resolution characterizes high-risk gastric cancer precursors. NPJ Precis. Oncol. 9:52.

Ibrahim MM. 2010. Subcutaneous and visceral adipose tissue: structural and functional differences. Obes. Metab. 7:64–65.

Inoue T et al. 2022. Non-invasive human skin transcriptome analysis using mRNA in skin surface lipids. Commun. Biol. 5:215.

Jaeger-Ruckstuhl CA et al. 2024. Signaling via a CD27-TRAF2-SHP-1 axis during naive T cell activation promotes memory-associated gene regulatory networks. Immunity. 57:287–302.e12.

Jin L et al. 2022. Lactate receptor HCAR1 regulates cell growth, metastasis and maintenance of cancerLspecific energy metabolism in breast cancer cells. Mol. Med. Rep. 26. doi: 10.3892/mmr.2022.12784.

Kaur TP, Verma R, Choudhary RK. 2021. Introduction to mammary gland and its cell types. In: Stem Cells in Veterinary Science. Springer Nature Singapore: Singapore pp. 25–37.

Kawasaki K. 2018. The Origin and Early Evolution of SCPP Genes and Tissue Mineralization in Vertebrates. In: Biomineralization. Springer Singapore pp. 157–164.

Khan S, Fitch S, Knox S, Arora R. 2022. Exocrine gland structure-function relationships. Development. 149. doi: 10.1242/dev.197657.

Kolberg L, Raudvere U, Kuzmin I, Vilo J, Peterson H. 2020. gprofiler2 -- an R package for gene list functional enrichment analysis and namespace conversion toolset g:Profiler. F1000Res. 9:709.

Ku GM et al. 2012. Research resource: RNA-Seq reveals unique features of the pancreatic β-cell transcriptome. Mol. Endocrinol. 26:1783–1792.

Kumegawa K et al. 2022. GRHL2 motif is associated with intratumor heterogeneity of cis-regulatory elements in luminal breast cancer. NPJ Breast Cancer. 8:70.

Lazarescu O et al. 2025. Human subcutaneous and visceral adipocyte atlases uncover classical and nonclassical adipocytes and depot-specific patterns. Nat. Genet. 57:413–426.

Li CM-C et al. 2020. Aging-associated alterations in mammary epithelia and stroma revealed by single-cell RNA sequencing. Cell Rep. 33:108566.

Li D et al. 2018. OXTR overexpression leads to abnormal mammary gland development in mice. J. Endocrinol. 239:121–136.

Li D et al. 2021. Oxytocin receptor induces mammary tumorigenesis through prolactin/p-STAT5 pathway. Cell Death Dis. 12:588.

Liu H, Gruber CW, Alewood PF, Möller A, Muttenthaler M. 2020. The oxytocin receptor signalling system and breast cancer: a critical review. Oncogene. 39:5917–5932.

Lycke N, Bemark M, Spencer J. 2015. Mucosal B Cell Differentiation and Regulation. In: Mucosal Immunology. Elsevier pp. 701–719.

McManaman JL, Reyland ME, Thrower EC. 2006. Secretion and fluid transport mechanisms in the mammary gland: comparisons with the exocrine pancreas and the salivary gland. J. Mammary Gland Biol. Neoplasia. 11:249–268.

Medina D. 1996. The mammary gland: a unique organ for the study of development and tumorigenesis. J. Mammary Gland Biol. Neoplasia. 1:5–19.

Mohammad Nezhady MA, Modaresinejad M, Zia A, Chemtob S. 2023. Versatile lactate signaling via HCAR1: a multifaceted GPCR involved in many biological processes. Am. J. Physiol. Cell Physiol. 325:C1502–C1515.

Moll R, Divo M, Langbein L. 2008. The human keratins: biology and pathology. Histochem. Cell Biol. 129:705–733.

Montalbetti N, Dalghi MG, Albrecht C, Hediger MA. 2014. Nutrient transport in the mammary gland: calcium, trace minerals and water soluble vitamins. J. Mammary Gland Biol. Neoplasia. 19:73–90.

Nickell WB, Skelton J. 2005. Breast fat and fallacies: more than 100 years of anatomical fantasy. J. Hum. Lact. 21:126–130.

Oftedal OT, Dhouailly D. 2013. Evo-devo of the mammary gland. J. Mammary Gland Biol. Neoplasia. 18:105–120.

Öling S et al. 2024. A human stomach cell type transcriptome atlas. BMC Biol. 22:36.

Peaker M. 2002. The mammary gland in mammalian evolution: a brief commentary on some of the concepts. J. Mammary Gland Biol. Neoplasia. 7:347–353.

Pia-Foschini M, Reis-Filho JS, Eusebi V, Lakhani SR. 2003. Salivary gland-like tumours of the breast: surgical and molecular pathology. J. Clin. Pathol. 56:497–506.

Picardo M, Ottaviani M, Camera E, Mastrofrancesco A. 2009. Sebaceous gland lipids. Dermatoendocrinol. 1:68–71.

Pires ND, Dolan L. 2012. Morphological evolution in land plants: new designs with old genes. Philos. Trans. R. Soc. Lond. B Biol. Sci. 367:508–518.

Pond CM. 1977. The significance of lactation in the evolution of mammals. Evolution. 31:177.

Rauner G, Kuperwasser C. 2021. Microenvironmental control of cell fate decisions in mammary gland development and cancer. Dev. Cell. 56:1875–1883.

Reed AD et al. 2024. A single-cell atlas enables mapping of homeostatic cellular shifts in the adult human breast. Nat. Genet. 56:652–662.

Resseguie M et al. 2007. Phosphatidylethanolamine N-methyltransferase (PEMT) gene expression is induced by estrogen in human and mouse primary hepatocytes. FASEB J. 21:2622–2632.

Reversi A, Cassoni P, Chini B. 2005. Oxytocin receptor signaling in myoepithelial and cancer cells. J. Mammary Gland Biol. Neoplasia. 10:221–229.

Ritchie ME et al. 2015. limma powers differential expression analyses for RNA-sequencing and microarray studies. Nucleic Acids Res. 43:e47.

Robinson MD, McCarthy DJ, Smyth GK. 2010. edgeR: a Bioconductor package for differential expression analysis of digital gene expression data. Bioinformatics. 26:139–140.

Rosin FM, Kramer EM. 2009. Old dogs, new tricks: regulatory evolution in conserved genetic modules leads to novel morphologies in plants. Dev. Biol. 332:25–35.

Saitou M et al. 2020. Functional Specialization of Human Salivary Glands and Origins of Proteins Intrinsic to Human Saliva. Cell Rep. 33:108402.

Sharp JA et al. 2014. Bioactive functions of milk proteins: A comparative genomics approach. J. Mammary Gland Biol. Neoplasia. 19:289–302.

Slepicka PF, Somasundara AVH, Dos Santos CO. 2021. The molecular basis of mammary gland development and epithelial differentiation. Semin. Cell Dev. Biol. 114:93–112.

Sun W et al. 2024. OCA-B/Pou2af1 is sufficient to promote CD4+ T cell memory and prospectively identifies memory precursors. Proc. Natl. Acad. Sci. U. S. A. 121:e2309153121.

Sun Y et al. 2024. Restoring BARX2 in OSCC reverses partial EMT and suppresses metastasis through miR-186-5p/miR-378a-3p-dependent SERPINE2 inhibition. Oncogene. 43:1941–1954.

Tharmapalan P, Mahendralingam M, Berman HK, Khokha R. 2019. Mammary stem cells and progenitors: targeting the roots of breast cancer for prevention. EMBO J. 38:e100852.

Twigger A-J et al. 2022. Transcriptional changes in the mammary gland during lactation revealed by single cell sequencing of cells from human milk. Nat. Commun. 13:562.

Uebbing S et al. 2024. Evolutionary innovations in conserved regulatory elements associate with developmental genes in mammals. Mol. Biol. Evol. 41. doi: 10.1093/molbev/msae199.

Wagner W, Kania KD, Ciszewski WM. 2017. Stimulation of lactate receptor (HCAR1) affects cellular DNA repair capacity. DNA Repair (Amst.). 52:49–58.

Wang S, Sekiguchi R, Daley WP, Yamada KM. 2017. Patterned cell and matrix dynamics in branching morphogenesis. J. Cell Biol. 216:559–570.

Wang Z et al. 2023. GRHL2-controlled gene expression networks in luminal breast cancer. Cell Commun. Signal. 21:15.

Xu Y et al. 2017. Twist1 promotes breast cancer invasion and metastasis by silencing Foxa1 expression. Oncogene. 36:1157–1166.

Zhang H, Gao M, Zhao W, Yu L. 2023. The chromatin architectural regulator SND1 mediates metastasis in triple-negative breast cancer by promoting CDH1 gene methylation. Breast Cancer Res. 25:129.

