## Supplementary material for "Comparative Transcriptomics of Human Breast Tissue Suggests Conserved Epithelial Secretory Programs and Tissue-Associated Regulatory Specialization": SFigures

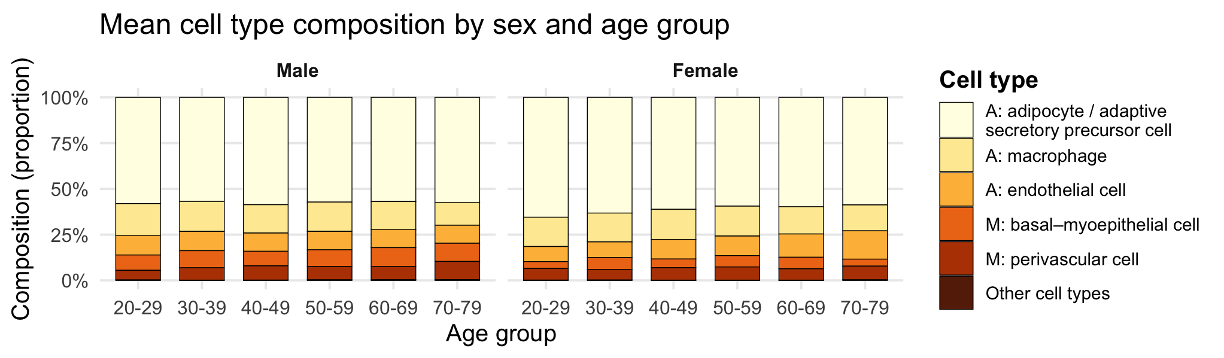

**Figure S1. Mean cell type composition in breast tissue by sex and age group**

Bulk RNA-seq data from GTEx breast tissue were deconvoluted using single-cell reference expression profiles to estimate relative cell-type composition. Stacked bar plots show mean fractional compositions averaged within each sex (male, female) and age group (10-year bins), with the y-axis normalized to 100%. Colors denote inferred cell populations: adipocyte / adaptive secretory precursor cell (A: Adipose), macrophages (A: Adipose), endothelial cells (A: Adipose), basal–myoepithelial cells (M: Mammary),perivascular cells (M: Mammary), and minor fractions classified as “Other cell types.” Single-cell RNA-seq reference datasets of  breast tissue and subcutaneous adipose were downloaded at CELLxGENE Discover portal (CZI Cell Science Program et al. 2025).

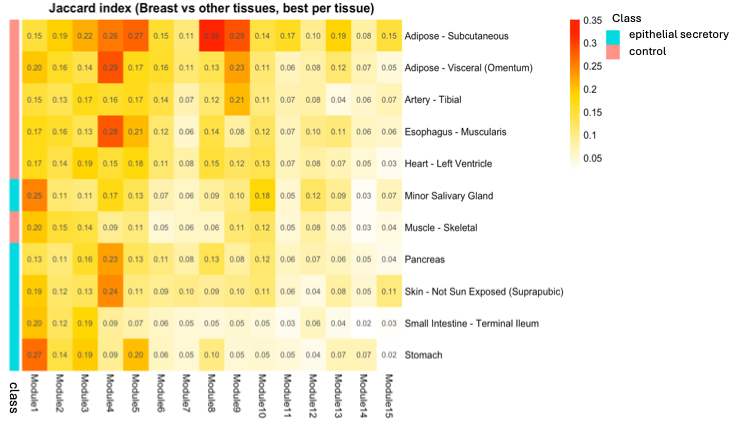

**Figure S2. Cross-tissue overlap of breast co-expression modules.**

Heatmap showing Jaccard indices between breast tissue co-expression modules and modules derived from other GTEx tissues. For each comparison tissue, the Jaccard index represents the maximum overlap observed between a given breast module and any module from that tissue (“best-per-tissue” overlap). Rows correspond to comparison tissues, and columns correspond to breast co-expression modules. Tissues are annotated as tissues containing epithelial secretory components or non-glandular tissues based on the tissue grouping used in this study. Warmer colors indicate higher Jaccard similarity.

CZI Cell Science Program et al. 2025. CZ CELLxGENE Discover: a single-cell data platform for scalable exploration, analysis and modeling of aggregated data. Nucleic Acids Res. 53:D886–D900.
